# TREM2 R47H exacerbates immune response in Alzheimer’s disease brain

**DOI:** 10.1101/499319

**Authors:** Olena Korvatska, Kostantin Kiianitsa, Alexander Ratushny, Mark Matsushita, Neal Beeman, Wei-Ming Chien, J-I Satoh, Michael O. Dorschner, C. Dirk Keene, Theo K. Bammler, Thomas D. Bird, Wendy H. Raskind

## Abstract

The R47H variant in the microglial TREM2 receptor is a strong risk factor for Alzheimer’s disease (AD). To characterize processes affected by R47H we performed integrative network analysis of genes expressed in brains of AD patients with R47H, sporadic AD without the variant and patients with polycystic lipomembranous osteodysplasia with sclerosing leukoencephalopathy (PLOSL), a systemic disease with early onset dementia caused by loss-of function mutations in TREM2 or its adaptor TYROBP. While sporadic AD had few perturbed microglial and immune genes, TREM2 R47H AD demonstrated upregulation of interferon type I response and pro-inflammatory cytokines accompanied by induction of NKG2D stress ligands. In contrast, PLOSL had distinct sets of highly perturbed immune and microglial genes that included inflammatory mediators, immune signaling, cell adhesion and phagocytosis. TREM2 knock-out in THP1, a human myeloid cell line that constitutively expresses the TREM2-TYROBP receptor, inhibited response to the viral RNA mimetic poly(I:C), and overexpression of ectopic TREM2 restored the response. Compared to wild type protein, R47H TREM2 had higher stimulatory effect on the interferon type I response signature. Our findings point to a role of the TREM2 receptor in the control of the interferon type I response in myeloid cells and provide insight regarding the contribution of R47H TREM2 to AD pathology.

## INTRODUCTION

A proactive role of brain immune cells, microglia, in the pathogenesis of Alzheimer’s disease (AD) gained recognition after discovery of AD-associated variants in the Triggering Receptor Expressed on Myeloid Cells 2 (*TREM2*) gene that is only expressed on immune cells of myeloid origin. Further support came from large-scale GWAS that identified multiple additional immune risk factors (OR 0.9-1.15) (1). Many of these immune molecules have either microglia-specific expression (*e*.*g*., CD33, the MS4As, INPP5D, SPI1) or are enriched in microglia (*e*.*g*., ABCA7, CR1, CLU). Of all microglial genetic AD risk factors, TREM2 R47H variant confers the highest risk of disease development, with a risk similar to that of APOEε4 (OR 4.5-2.6) (2, 3). Loss-of-function mutations in the TREM2 receptor or its adaptor, TYROBP, cause polycystic lipomembranous osteodysplasia with sclerosing leukoencephalopathy (PLOSL; OMIM # 221770), a recessive systemic disorder that presents with early onset dementia (4) or familial frontotemporal dementia (5).

TREM2 is involved in multiple aspects of myeloid cell function and interaction with other cell types (6). In microglia, it regulates cell chemotaxis (7), enhances phagocytosis (8, 9) and influences microglia survival, proliferation and differentiation (10). The complexity of TREM2 functioning is revealed in cell/tissue content dependent responses to stimuli of different nature, as well as their timing. For instance, whereas isolated TREM2-deficient cells increased production of pro-inflammatory molecules *in vitro* (8, 11), studies of intact tissues demonstrated that both pro- and anti-inflammatory roles of TREM2 in brain immunity are dependent on timing after stimulation (12), stage of pathology (13) and genetic background (14). In addition, a soluble form of TREM2, a product of cleavage or alternative splicing, may have a separate function as an extracellular signaling molecule that promotes cell survival (15, 16). TREM2 binds lipids and lipoproteins including the known AD risk factors ApoE and ApoJ/CLU (17-19) and mediates myelin clearance (20). Also, TREM2 binds amyloid beta peptide activating microglia cytokine production and degradation of internalized peptide (9).

By exome sequencing of families affected with late onset AD we identified multiple carriers of the TREM2 R47H variant (21). TREM2 R47H AD patients demonstrated a shortened course of disease, pronounced synucleinopathy, changed microglial morphology and decreased level of the microglial marker Iba1, suggesting that compromised TREM2 signaling has a strong effect on microglia function in the disease. Herein, we performed RNA expression profiling of brain tissues from subjects with TREM2 R47H AD, sporadic AD without the variant (sAD) and normal aged-matched controls to assess the effect of R47H variant on brain immune gene networks. We also analyzed brain tissues of PLOSL patients with bi-allelic loss-of-function mutations in TREM2 or TYROBP. To further investigate the role of TREM2 and its R47H variant, we knocked out the gene in the THP1 cells, a human myeloid line that expresses a functional TREM2-TYROBP receptor.

## MATERIALS AND METHODS

### Subjects

#### Familial AD and frontotemporal dementia (FTD) patients with R47H

Through the University of Washington (UW) Alzheimer’s Disease Research Center, we acquired brain autopsy material of nine TREM2 R47H carriers from four unrelated families affected with AD or FTD (Tables S1, S2). Flash-frozen brain tissues from two R47H carriers were used for RNA-seq analyses. RNA isolated from formalin-fixed, paraffin-embedded (FFPE) sections of eight R47H carriers with AD were used for gene expression by Nanostring nCounter.

#### Sporadic AD and controls

Frozen and fixed autopsy tissues from neuropathologically confirmed AD cases and aged non-demented controls were obtained from the UW Neuropathology Core Brain Bank (Tables S1, S2).

#### PLOSL cases caused by loss-of-function mutations in TREM2-TYROBP receptor genes

Brain tissues were obtained from the UW Neuropathology Core Brain Bank and the Meiji Pharmaceutical University (MPU), Tokyo, Japan. One PLOSL patient was homozygous for a missense variant D134G in *TREM2* (4), another patient was homozygous for a splice site mutation, c.482+2T>C, in *TREM2* (22) and two were homozygous for c.141delG in *TYROBP*(23) (Tables S1, S2). FFPE sections were used for RNA isolation and Nanostring nCounter analysis.

Under protocols approved by the Institutional Review Boards of the University of Washington and the Meiji Pharmaceutical University, all subjects had previously given informed consent to share and study autopsy material. All methods for processing and analyzing the brain autopsy tissues followed relevant guidelines and regulations.

### Cell culture and cytokine stimulation

A human microglia cell line HMC3 and human myeloid cell lines THP1, U937 and MOLM13 were obtained from the ATCC. Cells were cultured in DMEM (HMC3) or RPMI (the others) supplemented with 10% fetal bovine serum. Pro-inflammatory stimulation with LPS (Sigma, #055:B5, 0.5 μg/ml) and IFNγ (PeproTech, #300-02, 150 U/ml) was performed for 24 hrs. Stimulations with IL-4 (PeproTech, #200-04, 20 ng/ml) or IL-10 (PeproTech, #200-10, 20 ng/ml) were similarly performed for 24 hrs.

### Genome editing of *TREM2* with CRISPR/Cas9

A THP1 derivative that stably expresses Cas9 (THP1-CAS9) was generated using a lentiviral construct (Addgene, lentiCas9-Blast, #52962) after blasticidine selection. sgRNA for targeting *TREM2* were selected using the online CRISPR design tool (crispr.mit.edu) (24). THP1-CAS9 cells were nucleofected with sgRNA using a protocol recommended for THP1 (Lonza, Amaxa 4D Nucleofector, Protocol #292). Cutting efficiency for each guide was measured using GeneArt Genomic Cleavage Detection Kit (ThermoFisher, #A24372). Based on % indel efficiency, sgRNA within exon 2 (sense: ACTGGTAGAGACCCGCATCA) was chosen for subsequent experiments. TREM2-negative cells were enriched by cell sorting after immunostaining of live cells. Colonies grown from single sorted cells were sequenced; clones with frame-shifting indel mutations inactivating all *TREM2* alleles were tested for absence of protein by Western blotting.

### TREM2 cloning and overexpression

Coding sequences of human *TREM2* (NM_018965.3), common variant (CV) and R47H variant, were synthesized and cloned into doxycycline-inducible lentiviral pCW57-MCS1-2A-MCS2 vector (Addgene, #71782). THP1 TREM2 KO cells were transduced with lentiviral particles expressing either of wild-type TREM2, R47H TREM2 or GFP proteins (Addgene; pCW57-GFP-2A-MCS, #71783). After 2-weeks of puromycin selection, resistant cells were induced with doxycycline (100 ng/ml) and tested for TREM2 and GFP expression by qRT-PCR and by FACS analyses.

### THP1 differentiation into macrophages and stimulation of interferon type I response

THP1 differentiation to macrophages was performed using 5ng/ml phorbol 12-myristate 13-acetate (PMA) for 48 hrs followed by 24 hrs of recovery culturing without PMA (25, 26). IFN I response was induced by high molecular weight poly(I:C) complexes with LyoVec transfection reagent (Invivogen, tlrl-piclv, 500 ng/ml) or with a combination of poly(I:C)/LyoVec and interferon beta (PeproTech, #300-02, 100Units/ml) for 24 hrs. When indicated, TREM2 or GFP expression was induced by 100 ng/ml doxycycline 16-18 hr prior to the IFN I response stimulation.

### qRT-PCR analysis

Total RNA from harvested cells was isolated with RNAeasy kit (Qiagen; #74106); cDNA was synthesized with SensiFast (Bioline, BIO-65053). TaqMan® Fast Advanced Master Mix and the following TaqMan Gene Expression Assays (ThermoFisher, #4331182) were used for qRT-PCR: MICB: Hs00792952_m1, IFNB: Hs01077958_s1, IRF7: Hs01014809_g1, IFIH1: Hs00223420_m1 and reference gene TBP: Hs00427620_m1. Assays were performed on StepOnePlus real time PCR machine and analyzed with StepOne software (Applied Biosystems) using the 2(-Delta Delta C(T)) method (27). All measurements were performed in technical duplicates.

### Antibodies and immunostaining

Live cells were immunostained with fluorescent polyclonal antibodies to TREM2 (R&D Systems #MAB17291) and analyzed by FACS. For western blotting, 50 µg of whole cell lysates were resolved on polyacrylamide gel, transferred to PVDF membrane and stained with polyclonal TREM2 antibodies (Cell Signaling Technology, #91068) at 1:1000 dilution.

### Gene expression analyses

#### Whole transcriptome RNA-seq of brain tissues

Total RNA was isolated from flash-frozen brain tissues, and ribosomal RNA was depleted using the Ribo Zero Gold Magnetic system (Epicentre/Illumina, San Diego, CA). RNA-seq libraries were prepared with the ScriptSeq v2 kit (Epicentre) and subjected to paired-end sequencing (2 x 100 bp) on an Illumina HiSeq 2500 instrument. A mean read number of 6×10^7^ was generated per sample. The following data analysis tools were used: FastQC 0.9.6 for visualizing read QC https://www.bioinformatics.babraham.ac.uk/projects/fastqc; Bowtie 2.2.1 https://sourceforge.net/projects/bowtie-bio/files/bowtie2/2.2.1/ and TopHat 2.0.11 https://ccb.jhu.edu/software/tophat/index.shtml for read alignment and splice site discovery; Cufflinks 2.1.1 http://cole-trapnell-lab.github.io/cufflinks/releases/v2.1.1/ for transcript assembly and quantification; MISO 0.5.2 https://sbgrid.org/software/titles/miso for quantitating isoform expression levels; cummeRbund 2.0.0 http://compbio.mit.edu/cummeRbund and IGV 2.3.14 http://software.broadinstitute.org/software/igv/download for data visualization.

#### Gene expression analysis by nCounter system (NanoString Technologies)

Total RNA was isolated from FFPE samples. For each sample, four 10 μm thick FFPE sections were processed using the Recover All RNA isolation kit (Ambion, Termo Fisher) following the manufacturer’s protocol. RNA yield and fragment-length distribution were measured by Qubit Fluorimeter (Molecular Probes, Termo Fisher) and 2100 Bioanalyzer (Agilent Technologies). Typical yield was 0.8-1μg per 10 μm section. As expected, RNAs from archived FFPE samples were highly degraded (RIN 2-3), however more than 80% of RNA species were over 80 bp allowing analysis by Nanostring technology (hybridization with the 70 nucleotides gene-specific probe). Total RNA (1 μg) was used for hybridization on a NanoString nCounter platform according to the manufacturer’s instruction. Samples in which >50% genes had raw counts below the cutoff of 20 (average of 8 negative controls plus 2 standard deviations, 95% confidence interval) were excluded from analysis.

For gene expression analysis we used the PanCancer Immune panel, which contains 730 immunity-related and cancer genes together with 40 validated housekeeping genes, and a custom CodeSet containing 30 genes designed at Nanostring Technologies (NanoString Technologies). All probe sets were produced in a single batch. Data were normalized and log2 transformed using the nSolver 3.0 Analysis Software (NanoString Technologies). In brief, normalization was performed in two steps, 1) technical normalization using positive control spikes on each panel, and 2) input amount normalization using the mean expression of 30 housekeeping genes selected by a geNorm algorithm on the basis of stability of their expression through all experimental groups. All transcripts with raw counts below the threshold 20 (average of 8 negative controls plus 2 SD (∼ 95% confidence interval) were labeled as undetected. Only genes detected at a level of 30 or more row counts in at least 50% of our samples were included in the analysis (N=493).

### Gene set and pathway analyses

Gene Set Analysis (GSA) (28) was used to calculate significant enrichment of known transcriptional signatures in gene expression data from RNA-seq. We used 10,295 gene sets from the Molecular Signature Database version 4.0(29). False discovery rate and *P-*value estimates were computed for each gene set based on 1000 separate permutation distributions.

To identify biologically relevant pathways in differentially expressed genes from Nanosting nCounter data, we used Ingenuity Pathway Analysis (IPA, Ingenuity Systems, Qiagen) and Metascape (metascape.org). IPA queries a proprietary knowledge database and calculates significance of genes/pathways enrichment using Fisher’s exact test. Metascape queries public databases (Reactome, MSigDB, GO, KEGG Pathway, CORUM), and enrichment is calculated by hypergeometric test. Additionally, Metascape performs analysis of protein-protein interactions (PPI, data from BioGrid database) in which subnetworks with high local network connectivity are found using the Molecular Complex Detection (MCODE) algorithm (30). 800 genes on the Nanostring panel were used as a baseline for enrichment calculations. Visualization of protein-protein association networks that include known protein interaction and functional connection was done using STRING database (http:/string-db.org) (31).

### Statistical analysis

Hypergeometric distributions were evaluated using the online calculator https://systems.crump.ucla.edu/hypergeometric/index.php. Other statistical tests were performed using GraphPad Prism software.

## RESULTS

### Nanostring profiling reveals perturbed immune, microglial and AD-related genes in TREM2 R47H AD and few alterations in sAD

Hippocampi of TREM2 R47H AD patients (N=8) were compared with sAD (N=15) and normal aged controls (N=17); the latter two groups were chosen to minimize covariation with age, gender, disease stage, post-mortem interval and RNA quality (Tables S1, S2). Hippocampus is one of the most affected brain regions in AD, and in a previous study we observed the most pronounced changes in microglial markers in hippocampi of R47H carriers (21). We used Nanostring nCounter, a mid-throughput nucleic acid hybridization-based platform suitable for analysis of RNA from formalin-fixed paraffin-embedded (FFPE) material. A predesigned PanCancer Immune panel (770 genes) contained the main human immunity-related pathways; of those, 156 genes were microglia-specific, i.e., expressed at least 10-fold higher in microglia than in other brain cell populations (32). Of 800 genes on the panel, 551 had a detectable expression in FFPE samples and 493 of these, including 134 microglia-specific genes, were included in analysis.

At present, Nanostring nCounter is the method of choice for profiling RNA from FFPE-archived tissues. A recent study of FFPE AD brains found no major effect of post-mortem interval and RNA degradation on gene expression data (33). Also, this and other studies reported high concordance of Nanostring expression data in frozen and FFPE material from the same case (34), and we confirmed this for our samples and experimental setting. RNA isolated from flash frozen or FFPE tissue of the same individual showed high Pearson’s correlation of gene expression (r^2^ = 0.84). Additionally, the robustness of the assay was tested in the “discovery mode” using RNA from FFPE hippocampi of patients with X-linked Parkinsonism with Spasticity (XPDS; OMIM# 300911; N=2), a monogenic disorder caused by a variant in the *ATP6AP2* gene (21). The mutation alters ATP6AP2 splicing resulting in the abnormally high expression of the isoform that skips exon 4 (e3-e5). We found that e3-e5 was significantly increased in XPDS brains, as previously shown on RNA prepared from fresh blood cells of carriers (Figure S1). Thus, even in a small group of FFPE samples, Nanostring nCounter reliably detects gene expression changes caused by a mutation of predicted effect.

The TREM2 R47H AD group had the largest number of differentially expressed genes (104 vs 20 in sAD, FDR<0.1, Figure 1A, B). 75% of differentially expressed genes in sAD were shared with TREM2 R47H AD (Figure 1C). We measured the relative abundance of microglial cells in the brain tissue by scoring expression of 134 microglia-specific genes (MG signature, Figure 1D). Both AD groups showed similar increase in microglia cellularity compared to controls, suggesting that the observed increase in immune genes in TREM2 R47 AD is not due to increased number of microglia.

**Figure 1.**
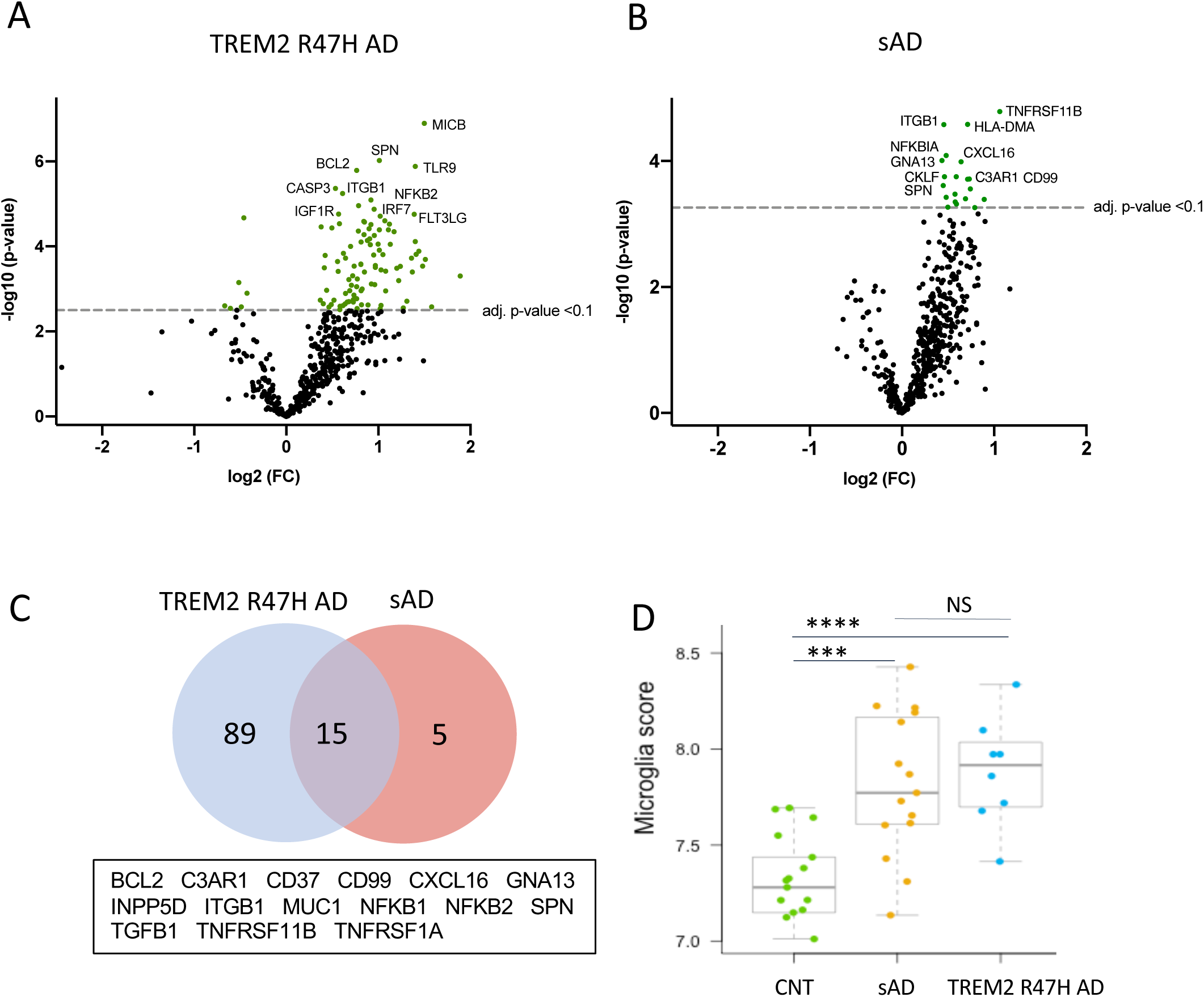
Differentially expressed (DE) genes in hippocampi of TREM2 R47H AD and sAD patients. (A-D) Volcano plots displaying DE genes in TREM2 R47H AD (A) and sAD (B) against a baseline of gene expression in controls. The y-axis corresponds to the log_10_(p-value), and the x-axis displays the log_2_ (FC, fold changes) value. (C). Venn diagram and the list of shared DE genes (FDR < 0.1) between TREM2 R47H AD and sAD. (D) Microglia cell scores were calculated by nSolver (v.3) as an average log-transformed expression of the 134 microglia-specific genes that allow comparison of cell abundance across samples (71). Each unit increase in a cell score calculated from log2 transformed data corresponds to a doubling of microglia abundance. (NS - non-significant; *** - p-value < 0.001, **** - p-value < 0.0001, one-way ANOVA, Tukey’s multiple comparison test).

We also assessed whether TREM2 R47H affects transcript levels of TREM2 itself, as well as other known AD genes and risk factors. Five of 16 genes with measurable expression were perturbed in TREM2 R47H AD while none reached significance in sAD (Figure 2). Four genes, including TREM2, were upregulated and PSEN2 was downregulated. Thus, the R47H variant has a profound effect on expression of AD risk factors.

**Figure 2.**
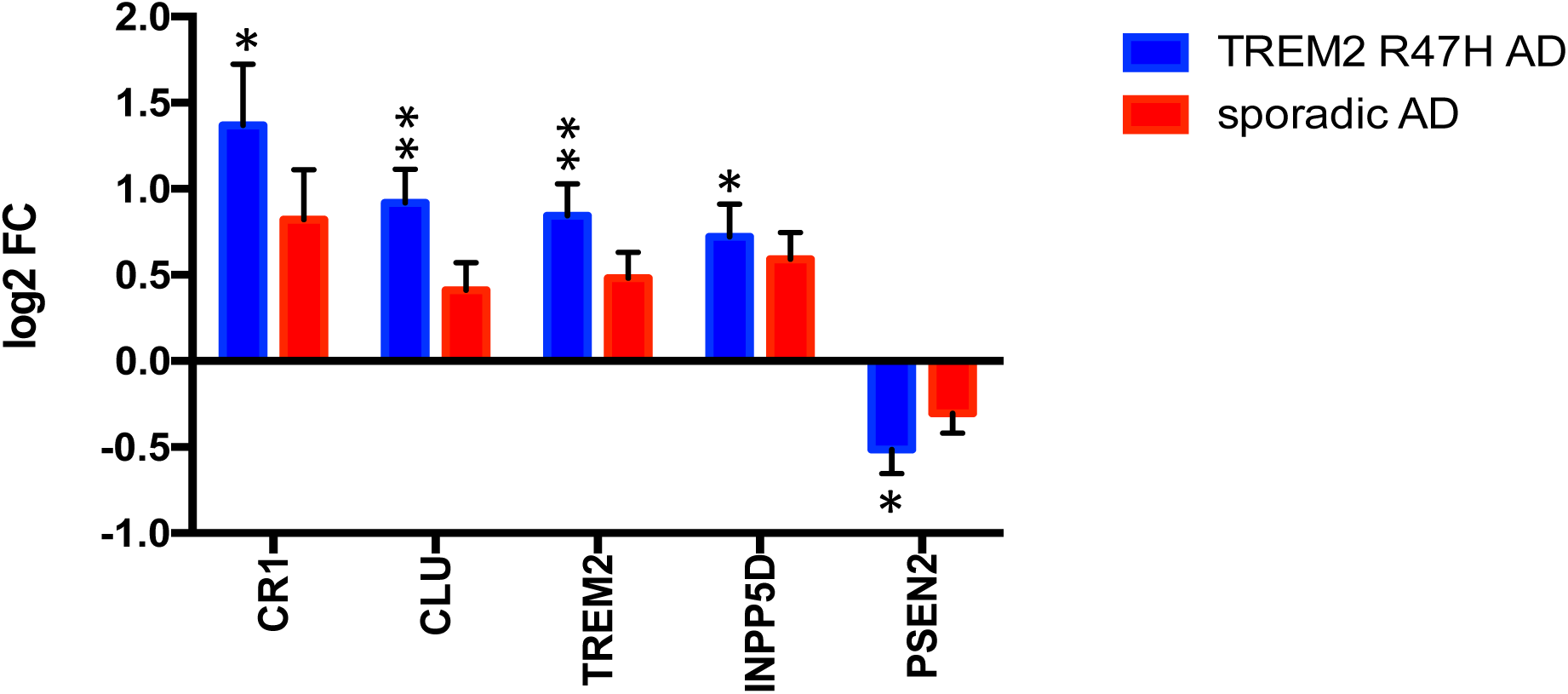
Perturbed expression of AD genes/risk factors in TREM2 R47H AD. AD-associated genes altered (FDR < 0.1) in at least one condition against a baseline of gene expression in controls are shown. Data are presented as mean ± SD, (* - adj p-value < 0.05; ** - adj p-value < 0.01); y-axis displays log2 (FC, fold changes) value.

### Pro-inflammatory immune networks and pathways are activated in TREM2 R47H AD brains

The most perturbed pathways in TREM2 R47H AD patients comprised “WNT/Ca^++^ Signaling”, “Role of RIG1-like Receptors in Antiviral Innate Immunity”, “Inflammasome Pathway” and “Neuroinflammation Signaling Pathway” (Table 1). Analysis of the protein-protein interaction (PPI) network revealed enriched GO terms corresponding to regulation of cytokine production and I-kappaB kinase/NF-kappaB signaling and identified three functional modules corresponding to type I interferon response, ligand-receptor interaction, and a module formed by PSEN2 and mediators of apoptosis (Figure 3). In contrast, sAD produced a heterogeneous group of overrepresented pathways driven by the TNF receptor superfamily members (TNFRSF11b and TNFRSF1A) and the NF-kB complex and its regulators (Table 1). These findings corroborated a meta-analysis of sAD transcriptomes that identified the TNF receptor superfamily genes, such as TNFRSF11B and TNFRSF1A, and components of NFKB signaling, such as NFKBIA, NFKB1 and RELA, among the top up-regulated pathways in sAD (35).

**Table 1.** Top Canonical Pathways from Ingenuity pathway analysis of DE genes in TREM2 R47H AD and sAD.

| **TREM2 R47H AD** |  |  |  |  |
| --- | --- | --- | --- | --- |
| **Ingenuity Canonical Pathways** | **-log(p-value)** | **zScore** | **Ratio** | **Genes** |
| Wnt/Ca<sup>+</sup> pathway | 3.17 | 2.83 | 0.67 | NFKB2, NFATC3, NFATC4, NFKB1, NFATC2, EP300, CREBBP, NFATC1 |
| Role of RIG1-like Receptors in Antiviral Innate Immunity | 2.73 | 3.00 | 0.50 | MAVS, NFKB2, TRAF2, NFKB1, IRF7, IFIH1, EP300, CREBBP, CASP8, TRAF6, IKBKE |
| Inflammasome pathway | 2.64 | 2.65 | 0.64 | TLR4, NFKB2, NFKB1, NOD2, CASP1, CASP8, PYCARD |
| Neuroinflammation Signaling Pathway | 2.45 | 3.54 | 0.31 | TGFB1, NFATC2, TNFRSF1A, TICAM1, IL6R, TLR9, TREM2, CSF1R, TLR4, IRF7, CREBBP, TRAF6, NFATC4, CCL3, NFKB1, JAK2, EP300, CASP1, HLA-DOB, APP, MFGE8, NFATC1, PYCARD, NFKB2, NFATC3, PSEN2, BCL2, TICAM2, CASP3, CD200, CASP8, IKBKE |
| **sAD** |  |  |  |  |
| **Ingenuity Canonical Pathways** | **-log(p-value)** | **zScore** | **Ratio** | **Genes** |
| Induction of Apoptosis by HIV1 | 3.74 | 0.82 | 0.25 | NFKB2, TNFRSF11B, NFKB1, TNFRSF1A, BCL2, NFKBIA |
| OX40 Signaling Pathway | 3.25 | 1.00 | 0.21 | NFKB2, NFKB1, HLA-DMA, HLA-DRB3, BCL2, NFKBIA |
| Ceramide Signaling | 3.14 | 1.34 | 0.25 | NFKB2, TNFRSF11B, NFKB1, TNFRSF1A, BCL2 |
| PI3K/AKT Signaling | 3.08 | 1.64 | 0.19 | NFKB2, NFKB1, INPP5D, ITGB1, BCL2, NFKBIA |

**Figure 3.**
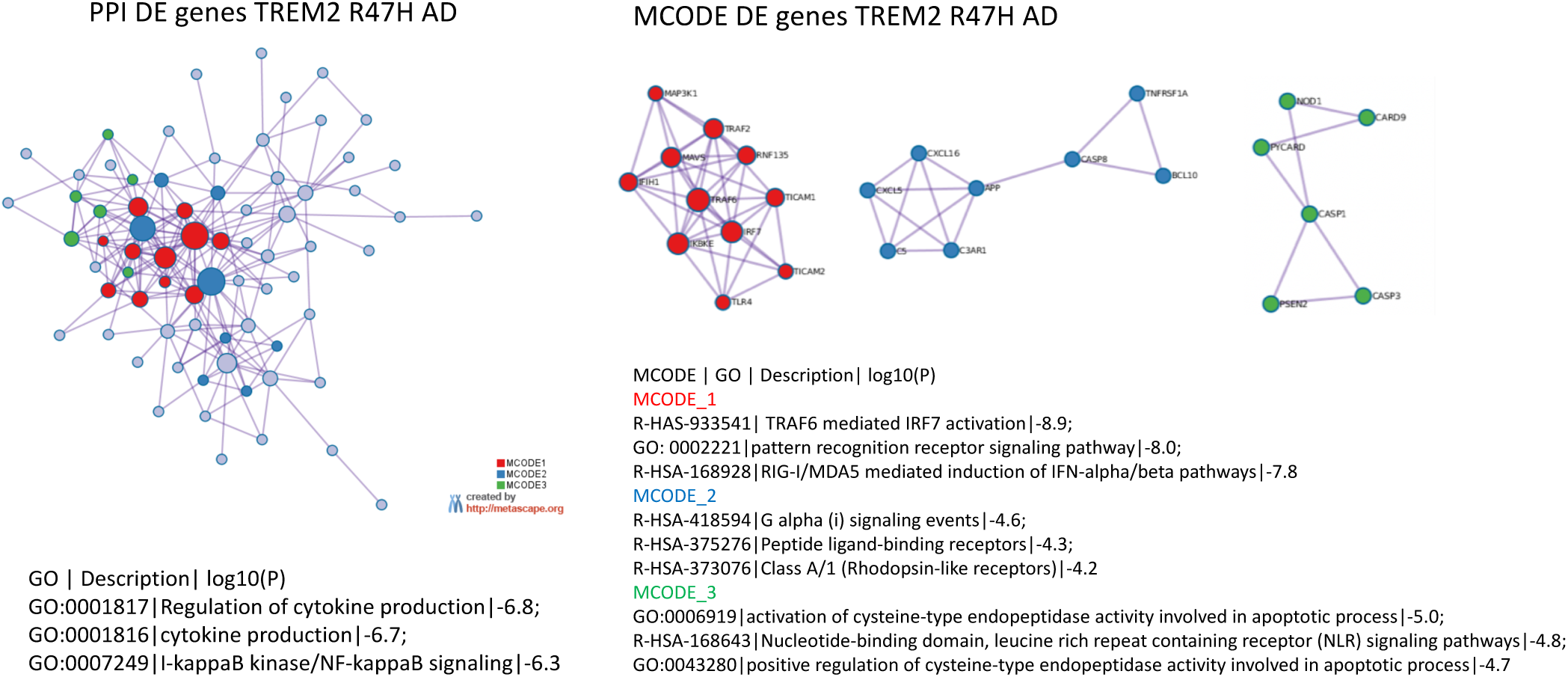
The protein interaction networks (PPI) and its tightly connected cores (MCODE) formed by differentially expressed genes in TREM2 R47H AD. PPI, MCODE and Gene Ontology (GO) term enrichment were determined using Metascape (www.metascape.org). 800 genes on the Nanostring panel were used as a background for enrichment calculations. Tables with the corresponding GO term enrichment are shown below. Column heading in table: GO - GO ID, Description: Representative GO term; Log10(P) - p-value for enrichment.

### TREM2 R47H AD brain manifests activation of antiviral response genes and “induced-self” NKG2D ligands

The activated antiviral interferon type I (IFN I) response and upregulated pro-inflammatory cytokines were the top immune signatures in TREM2 R47H brains (Table 1, Figure 4). We also identified additional upregulated genes known to act in the antiviral response (RNF135/Riplet, TLR9, TLR4 and BST2/tetherin) that were not part of the canonical IFN I response signature (Figure 4B). During IFN I response, multiple downstream interferon stimulated genes (ISG) are upregulated. We therefore queried a database of ISG (INTERFEROME) (36) and found that differentially expressed genes in TREM2 R47H AD were ISG enriched (hypergeometric test p-value is 0.03).

**Figure 4.**
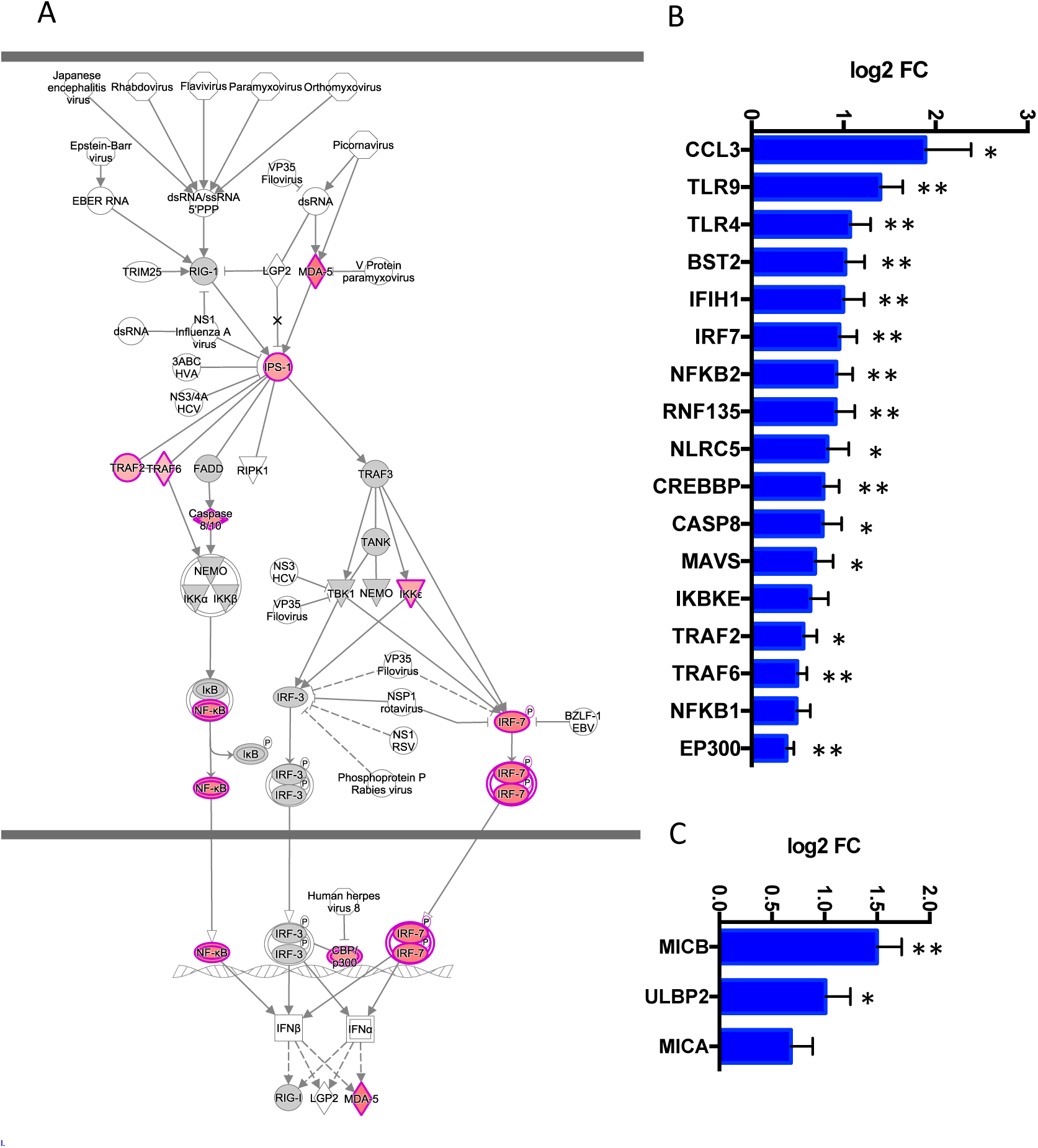
TREM2 R47H AD features upregulated Interferon type I response and MHC class I-like NKG2D ligands. (A) Role of RIG1-like receptors in antiviral innate immunity, IPA-generated diagram shows perturbed gene expression in TREM2 R47H AD brains. Red denote up-regulated genes, grey - not changed compared to control brains, white - not assessed. Perturbed expression (FDR < 0.1) of IFN I type response/antiviral genes (B) and MHC class I- like NKG2D ligands (C) Data are presented as mean ± SD (* - adj p-value < 0.05; ** - adj p-value < 0.01); y-axis displays log2 (FC, fold changes) value.

Dysregulated IFN I response activates multiple pro-inflammatory molecules causing cellular stress and cytotoxicity (37, 38). In TREM2 R47H AD brains, we observed upregulation of several MHC class I-like NKG2D ligands (MICB, MICA and ULBP2, Figure 4C). These molecules are known as “induced self” because their expression on the cell surface is induced by diverse stress agents, such as viruses, heat shock or oxidative damage. In the brain, MICB is expressed primarily by microglia (32), and it is the most significantly upregulated gene in TREM2 R47H AD (Figure 1A). We tested whether MICB may be upregulated in response to pro-inflammatory stimulation using a panel of human cell lines that share myeloid origin with microglia. Stimulation with a combination of LPS and IFNγ upregulated MICB expression in all tested cell lines, whereas stimulation with type 2 cytokines IL-4 or IL-10 had no effect (Figure S2).

### TREM2 R47H associated immune signatures are upregulated in brain transcriptomes of carriers

We also had flash-frozen unfixed hippocampus tissue suitable for RNA-Seq analysis from two of the TREM2 R47H carriers. To assess the top affected pathways at the transcriptome-wide level we sequenced total RNA of these two TREM2 R47H carriers and three normal age-matched controls. Table 2 shows the top 1% of differentially regulated gene sets in brains of TREM2 R47H carriers (23 of 2076 sets identified by Gene Set Analysis (GSA) (28). Of the 16 most upregulated sets 10 belonged to immune process/function, such as cytokine metabolism, antiviral response/TRAF6-mediated IRF7 activation and immune response. The second most upregulated group consisted of four signatures related to cell cycle/mitosis. Seven most down-regulated sets were related to ion and amino acid transport, endocytosis, glutamine metabolism and synaptic transmission (Table 2).

**Table 2.**
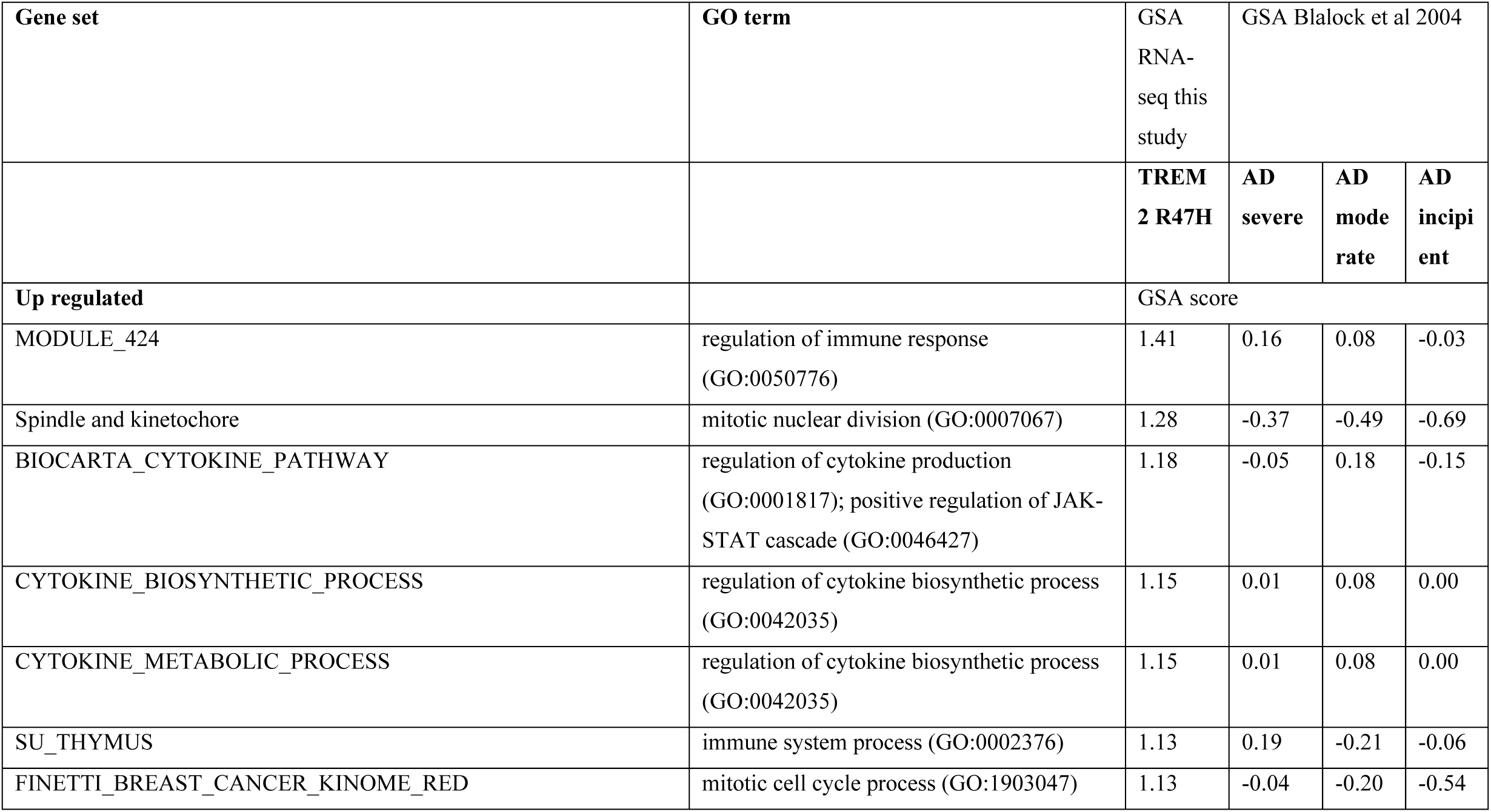
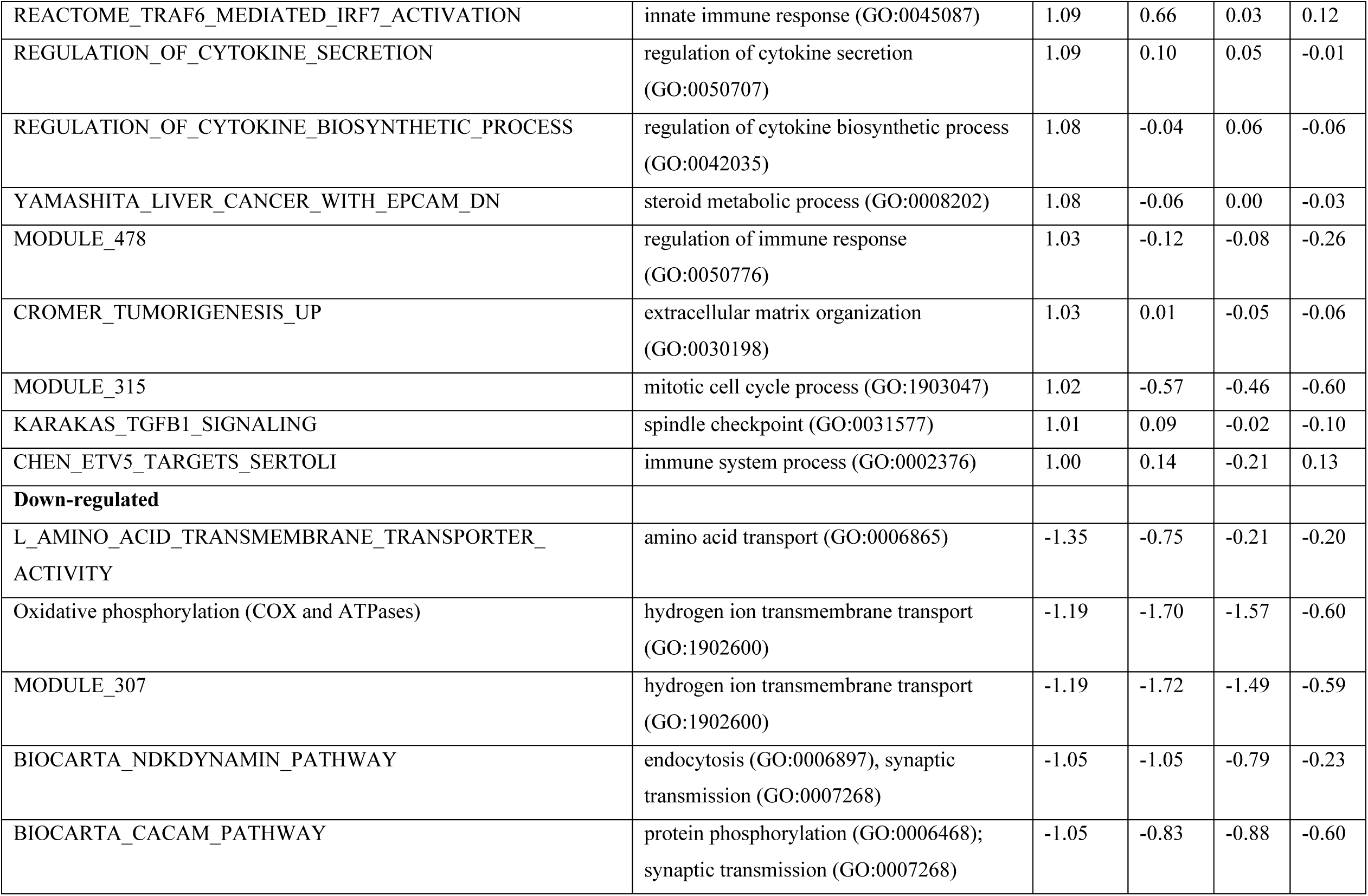
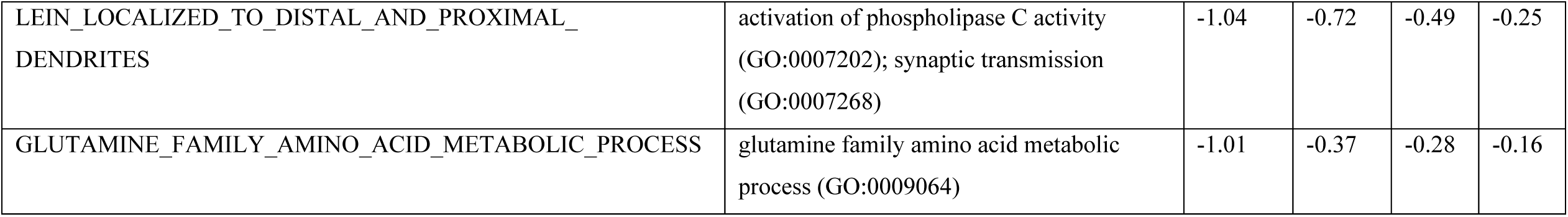
Top gene sets in TREM2 R47H brains are enriched by immunity-related GO terms. Shown are the top 1% of sets up/down regulated in carriers of TREM2 R47H, GSA scores ≥1 or ≤ −1 (GSA of RNA-seq data, this study). Enrichment of the same gene sets in sporadic AD was assessed separately by performing GSA of the publicly available dataset of the three sAD cohorts (GSA Blalock et al, 2004).

To distinguish whether the upregulation of immune genes was a characteristic of AD pathology (effect of disease) or due to abnormal function of the TREM2 immune receptor (effect of mutation) we performed GSA analysis on sAD data from the Blalock study (GDS810 (39)). This dataset has been used in several meta-analyses of differential gene expression in AD (40-43). The study compared hippocampi of the non-demented control group (N=9) with three progressive stages of AD: “incipient” (N=7), “moderate” (N=8) and “severe” (N=7). The TREM2 R47H and sAD groups shared a number of downregulated sets (Table 2), and their enrichment scores increased with disease progression. Notably, immune sets upregulated in the TREM2 R47H group were not enriched in sAD.

### Bi-allelic loss-of-function mutations in TREM2 or TYROBP cause gross perturbation of microglial networks in brains of PLOSL patients

PLOSL patients with bi-allelic mutations inactivating either subunit of the TREM2-TYROBP receptor develop an identical disease phenotype (4). This prompted us to compare changes in microglial and immune gene networks caused by the receptor complex loss with the effect of TREM2 R47H variant. We analyzed samples from four PLOSL patients with loss-of-function mutations in TREM2 (N=2) (4, 22) or TYROBP (N=2) (23) by Nanostring nCounter.

Differentially expressed genes in PLOSL (Figure 5A) were enriched for microglia-specific genes (40.4% vs 27% of expected distribution, hyper-geometric test, p = 0.027) consistent with the microglial origin of this disorder. In contrast, DE genes in TREM2 R47H AD were not MG enriched (30% vs 27% of expected distribution, hyper-geometric test, p = 0.21) suggesting that multiple cell types have contributed to the activation of immune response in this group. PLOSL and TREM2 R47H AD shared only 6 DE genes, less than 9 expected by chance, and no common pathways (Figure 5B-C). Top enriched pathways in PLOSL were “myeloid cell activation involved in immune response”; “bone development” and “Fc-gamma Receptor-mediated Phagocytosis”. We also compared fold changes of expression of all MG expressed in PLOSL and AD brains. PLOSL showed the highest magnitude of MG perturbation followed by TREM2 R47H AD (Figure 5D). All 19 differentially expressed MG in PLOSL formed a tight network orchestrated by the IRF8 transcription factor. The network was enriched with pro-inflammatory (HLA-DRA, IL18, TNFAIP3, ITGAX) and immune signaling molecules (SYK, JAK2, INPP5D, PTPRC), endo-phagocytosis mediators (C3, MSR1) and cell adhesion/motility receptors (CSF3R, ITGAX, OLR1, CD84, SELPLG, Figure 5E).

**Figure 5.**
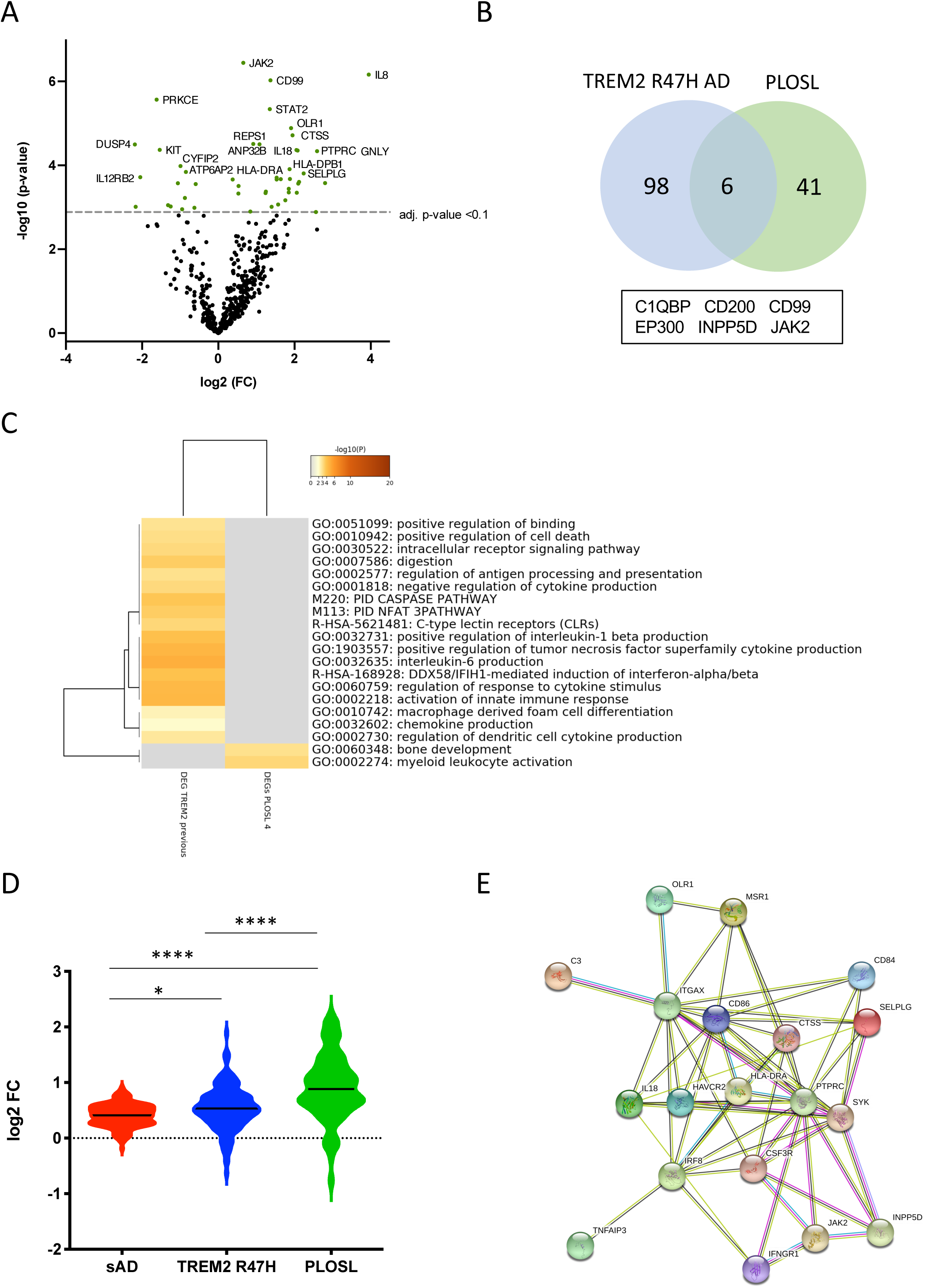
Differentially expressed and Microglia-specific genes in PLOSL. (A) Volcano plots displaying DE genes in PLOSL against a baseline of gene expression in controls. (B) Venn diagram and the list of shared DE genes (FDR < 0.1) between TREM2 R47H AD and PLOSL. (C) Enriched ontology clusters in DE TREM2 R47H AD and PLOSL. Metascape cluster analysis of GO term enrichment among DE genes (FDR < 0.1) between TREM2 R47H AD and PLOSL. 800 genes on the Nanostring panel were used as a background for enrichment calculations. GO ID, Description: Representative GO term; Log10(P) - p-value for enrichment. (D) Microglia-specific genes (MG) expression is highly perturbed in PLOSL as compared with AD groups. Distribution of fold changes in MG (N=134); X-axis displays log2 of fold changes between gene expression in cases vs controls. (* - p-value < 0.05; **** - p-value < 0.0001, one-way ANOVA, Tukey’s multiple comparison test); (E) Protein-protein association network formed by differentially expressed MG in PLOSL, as visualized by STRING database (http://string-db.org).

### TREM2 has a stimulatory role in the antiviral interferon type I response in myeloid cells

THP1 is a human monocytic cell line used to study monocyte-macrophage differentiation, signaling, innate immune and antiviral responses (44). It constitutively expresses the TREM2-TYROBP receptor and responds to known TREM2 stimulants, such as IL-4, similarly to primary macrophages (45). Using CRISPR/Cas9 genome editing, we generated TREM2 knockouts (KO) in THP1 (Figure S3) and created doxycycline-inducible derivatives that express common variant (CV) or R47H TREM2 on the KO background. To investigate the involvement of TREM2 in the IFN I response, THP1 derivatives were *in vitro* differentiated to macrophages with phorbol 12-myristate 13-acetate (PMA) (25, 26). IFN I response was induced by the high molecular weight RNA-mimetic poly(I:C) in a formulation that specifically activate the RIG-I/MDA5 (IFIH1) signaling pathway (47, 48). After 24 hr of stimulation, the response was evident from the upregulation of the interferon beta (IFNB) transcript (Figure 6A). Absence of TREM2 caused a notable decrease of IFNB stimulation in response to poly(I:C) that was restored by doxycycline-induced expression of ectopic CV or R47H TREM2 in the KO cells (Figure 6B). When normalized to the level of TREM2 expression, R47H had higher stimulatory effect on IFNB than CV TREM2 (Figure 6C). When derivatives were co-treated with interferon beta and poly(I:C), a combination known to potentiate the response, R47H TREM2 had superior effect on IFNB level, as well as on key IFN type I response genes, such as IRF7 and IFIH1 (Figure 7). In aggregate, our data indicate a previously unrecognized role of TREM2 as a positive regulator of IFN I response in myeloid cells expressing this receptor.

**Figure 6.**
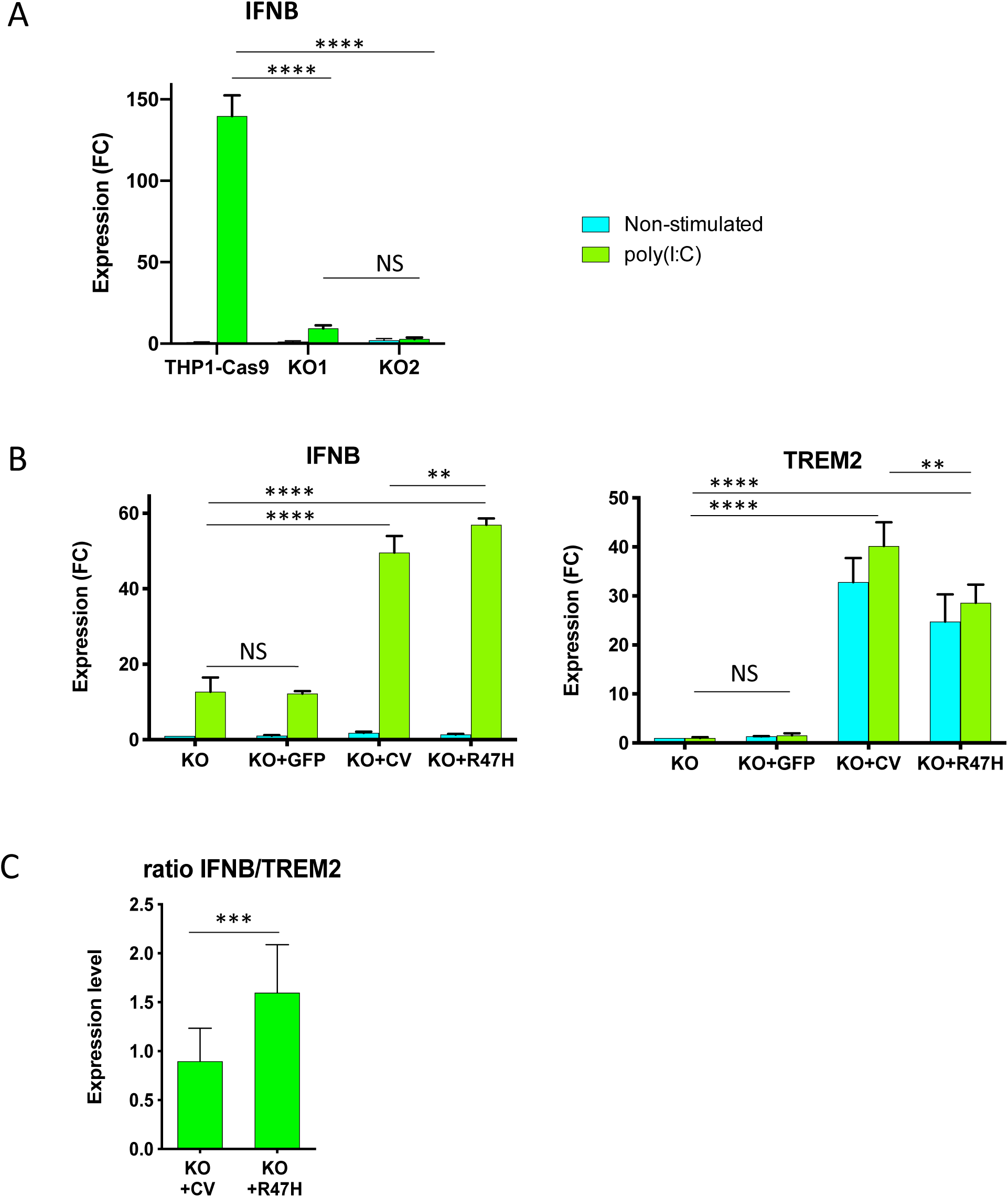
TREM2 stimulates IFN I response in THP1 cells. (A) Absence of TREM2 inhibited IFN I response in THP1 TREM2 KO. IFNB expression in non-stimulated cells and after 24h with poly(I:C) complexes was measured by qRT-PCR. THP1-Cas9, THP1 stably expressing Cas9 nuclease; KO1, KO2 are two independent TREM2 KO clones. IFNB expression was normalized to the level in unstimulated THP1-Cas9. FC – fold changes. (B) Overexpression of TREM2, but not GFP, restored the IFN I response. The assays were performed in biological triplicates and were repeated at least 2 times; A and B show representative experiments. KO - TREM2 KO1 cells; KO+GFP - TREM2 KO1 expressing GFP; KO+CV – TREM2 KO1 expressing common variant (CV) TREM2; KO+R47H - TREM2 KO1 expressing R47H TREM2. All cells were treated with doxycycline. TREM2 and IFNB expression was normalized to their levels in unstimulated KO. (C) Expression of R47H TREM2 induced higher level of IFNB as compared to CV TREM2. Ratio of IFNB to TREM2 expression were calculated for TREM2 KO1 THP1 expressing either CV TREM2 or R47H TREM2 after 24h poly(I:C) treatment from 8 independent biological replicates performed on different days. Data are presented as mean ± SD. Significance in A and B was calculated using two-way ANOVA, Tukey’s multiple comparison test; significance in C was calculated using unpaired two-tailed t-test. (NS – non-significant, ** - p < 0.01; *** - p<0.001; **** - p<0.0001).

**Figure 7.**
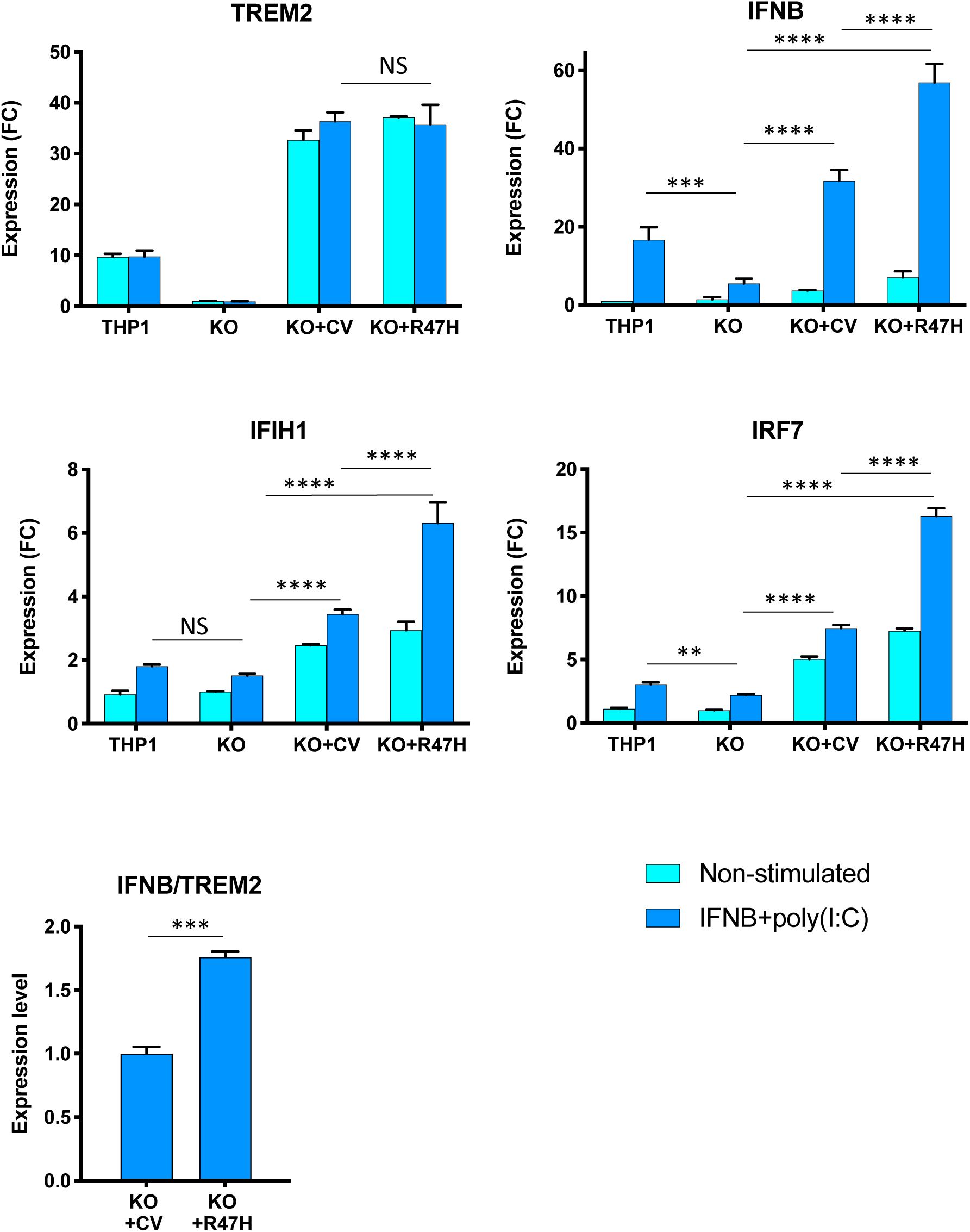
Effect of TREM2 on key IFN response genes upon stimulation with a combination of poly(I:C) and interferon beta. All cells were treated with doxycycline. Gene expression in non-stimulated cells and after 24 h stimulation was measured by qRT-PCR. THP1, unmodified cells; KO, TREM2 KO1 line; KO+CV, KO+R47H, TREM2 KO1 expressing common variant (CV) or R47H TREM2, respectively. Gene expression levels were normalized to expression in unstimulated KO. FC – fold changes. Assays were performed in biological triplicates and repeated 2 times; representative experiment is shown. Data are shown as mean ± SD (NS – non-significant, ** - p < 0.01; *** - p<0.001; **** - p<0.0001, two-way ANOVA, Tukey’s multiple comparison test). Ratio of IFNB to TREM2 expression were calculated for TREM2 KO1 THP1 expressing either CV TREM2 or R47H TREM2 after 24h poly(I:C) treatment from biological triplicates, data shown as mean ± SD (*** - p<0.001, unpaired two-tailed t-test).

## DISCUSSION

Pathogenic TREM2 variants of different effect are found in several neurodegenerative diseases. In PLOSL, mutations that inactivate TREM2, including premature stop codons (49, 50), splice site mutations (22, 51) and the missense variants D134G, K186N (4), Y38C, T66M, and V126G (5, 52, 53) have a recessive mode of inheritance. In contrast, AD-associated R47H has a dominant mode of inheritance. Whereas pathogenic missense PLOSL variants affect protein folding and stability decreasing TREM2 surface localization, R47H and other AD-associated mutations alter the ligand-binding interface of TREM2 (54, 55). Analysis of TREM2 R47H function in human and mouse reporter cells showed impaired recognition of certain anionic lipid ligands (20, 56). These findings are interpreted as weakened function not compensated by the normal allele (haploinsufficiency). Assuming this scenario, heterozygous carriers of known loss-of-function TREM2 variants in PLOSL families would also have increased AD susceptibility. To date, pedigrees with nine distinct TREM2-inactivating variants have been described across the world (57), and a study of AD burden in heterozygous mutation carriers in these PLOSL families would be required to test this hypothesis. Of several homozygous R47H individuals reported in Iceland none manifested a PLOSL-like early onset form of dementia, indicating that the variant protein has retained essential function (2).

We observed upregulation of immune and antiviral defense networks in the brain of heterozygous TREM2 R47H carriers who developed AD. It was exemplified by upregulation of pro-inflammatory cytokines and activation of IFN I response, an important line of defense against viruses. This pathway is activated upon sensing viral nucleic acids inside the cell triggering a cascade of immune responses that restrict viral propagation and eliminate infected cells. In the absence of viral infection, unchecked IFN I response is harmful, being a cause of Aicardi-Goutières syndrome (AGS), a group of monogenic interferonopathies that affect the CNS (58). A common RNA expression phenotype, an “interferon signature”, features upregulation of IFN I genes in AGS patients. There is no consensus as to the precise set of genes that constitute the signature (59), so the subsets may vary considerably between cases with different mutated genes that cause AGS and even between patients with the same mutation (60). Of note, some canonical IFN I response genes, such as STAT1, STAT2, ISG15, IFIT1 and IFIT2, were not perturbed in TREM2 R47H AD brains.

The IFN I response signature in the microglia is also a characteristic of brain aging in humans and mice (61). Pharmacologic blockage of IFN I signaling ameliorated age-related cognitive decline in mice. At present, it remains to be established which aging-associated processes may drive activation of the IFN I response in the brain. Under certain circumstances, the antiviral response may be triggered by reactivation of silenced endoretroviruses (ERV) or other transposable elements abundant in the human genome that mimic dsRNA produced during viral infection. For instance, DNA methyltransferase inhibition results in demethylation of silenced ERV, and their expression triggers a suicidal IFN I response (62, 63). In aging, transposable elements undergo profound epigenetic alterations, and their reactivation is observed in the context of organismal and cellular senescence, in human and other species (64). Inducing ERV activation in the mouse brain causes significant hippocampus-related memory impairment accompanied with robust activation of antiviral immune response mediated through Irf7; the pathology is attenuated in mice lacking cytosolic viral RNA sensor protein MAVS (65). The absence of TREM2 signaling in macrophages differentiated from KO-THP1 monocytes substantially blunted the response to viral dsRNA mimetic poly(I:C), seen as a decrease of IFNB induction. In the “add-back” experiment, induction of recombinant TREM2 restored IFNB production. Notably, the R47H variant induced greater IFNB increase than common variant (CV) protein indicating possible gain of function in the context of IFN I response stimulation. We propose a model in which TREM2 regulates the intensity of the antiviral IFN I response in specialized innate immune cells expressing this receptor, such as microglia and macrophages.

The epigenetic reprogramming that occurs in aging brain may activate transcription of silenced ERV or other transposable elements, *e*.*g*., through DNA demethylation or changes in histone modifications of their chromatin. Transcription from repeat elements positioned in sense and anti-sense orientation is able to generate endogenous dsRNA, which is recognized as foreign t and triggers chronic activation of the IFN I pathway. Trem2 is upregulated in the course of acute response to RNA virus in mouse macrophages (15), Trem2 and TREM2 levels increase with age in mouse and human brains (66), and TREM2 is elevated in the brain of R47H carriers. The proposed stimulatory role of the TREM2 risk allele in AD pathology is consistent with the small overlap of perturbed genes we observed in PLOSL and TREM2 R47H AD brains, as well as with inhibited IFN I response in THP1 TREM2 KO.

When this manuscript was in preparation, we became aware of another study that analyzed gene expression in TREM2 R47H brains. Prokop and colleagues studied several brain areas by IHC followed by Nanostring nCounter analysis of RNA isolated from FFPE sections (33). Notably, TREM2 R47H AD patients presented with a significant increase of senescent/dystrophic microglia in the hippocampus. R47H carriers also had elevated immune and microglial activation scores in frontal cortex and, to a lower extent, in hippocampus. Different choice of pre-designed gene panels in this study and in our work may in part explain the difference in the top observed immune gene signatures. The abundance of senescent microglia in TREM2 AD cases is an intriguing observation pointing to a possible role of this immune receptor in accelerated physiological aging of the microglia. In this light, our finding of co-occurrence of activated pro-inflammatory and IFN I pathways with upregulated NKG2D ligands, such as MICA/MICB and ULBP2, in TREM2 R47H AD brains may be viewed as part of complex senescence phenotype (67-69).

We acknowledge the limitations of our study engendered primarily by the rarity of autopsy samples from TREM2 R47H AD and PLOSL patients, most of which is limited to FFPE material. Notwithstanding these constraints, our findings in brains of R47H TREM2 carriers, together with demonstrated effects of TREM2 knockout and overexpression in myeloid cells, encourage further testing of the hypothesis that the variant exacerbates the neurodegenerative process in AD via chronic activation of the antiviral immune defense.

## Acknowledgements

We are grateful to Drs. Zoran Brkanac and John Neumaier for critical comments. This work is based on a previous pre-print article at bioRxiv (70).

## Author contributions

Conceptualization O.K., K.K., W.H.R., Methodology A.R., K.K., M.O.D., T.K.B., Investigation O.K., K.K., A.R., M.M., N.B., W-M.C. Resources J-I.S., C.D.K., T.D.B, Writing – original draft O.K., K.K., Writing – review and editing T.B.D., W.H.R., Visualization O.K., A.R., K.K., Supervision O.K., W.H.R.

---

**Table S1.** Clinical and neuropathological characteristics of cases and controls.

| Subject ID | Description | Gender | Age at death | Affected status | Braak | CERAD | Contributing pathology / affected region | Gene mutation | Transcript on profiling | Reference |
| --- | --- | --- | --- | --- | --- | --- | --- | --- | --- | --- |
| 458 | Non-affected R47H TREM2 | F | 87 | Non AD | III (B2) | C0 |  | TREM2 R47H | RNA-seq | (Korvatska et al., 2015) |
| 414 | TREM2 R47H AD | F | 76 | AD | VI (B3) | C1 |  | TREM2 R47H | RNA-seq, Nanostring |  |
| 5799 | TREM2 R47H AD | F | 64 | AD |  |  |  | TREM2 R47H | Nanostring |  |
| 661 | TREM2 R47H AD | M | 60 | AD | VI (B3) | C3 | LB, neocortical (diffuse) | TREM2 R47H | Nanostring |  |
| 5790 | TREM2 R47H AD | F | 85 | AD | VI (B3) | C2 |  | TREM2 R47H | Nanostring | (Korvatska et al., 2015) |
| 1205 | TREM2 R47H AD | F | 66 | AD | V (B3) | C3 |  | TREM2 R47H | Nanostring | (Korvatska et al., 2015) |
| 1462 | TREM2 R47H AD | F | 89 | AD | V (B3) | C3 | LB, amygdala predominant | TREM2 R47H | Nanostring | (Korvatska et al., 2015) |
| 1558 | TREM2 R47H AD | M | 67 | FTLD, AD |  | C2 | Tauopathy | TREM2 R47H | Nanostring |  |
| 4731 | TREM2 R47H AD | F | 81 |  |  |  |  | TREM2 R47H | Nanostring |  |
| 520 | sAD | F | 67 | AD | VI (B3) | C3 |  |  | Nanostring |  |
| 1291 | sAD | M | 67 | AD | VI (B3) | C3 |  |  | Nanostring |  |
| 655 | sAD | M | 63 | AD | VI (B3) | C3 | LB, amygdala predominant |  | Nanostring |  |
| 297 | sAD | F | 62 | AD | VI (B3) | C3 |  |  | Nanostring |  |
| 5918 | sAD | F | 81 | AD | VI (B3) | C3 |  |  | Nanostring |  |
| 5964 | sAD | F | 85 | AD | VI (B3) | C2 |  |  | Nanostring |  |
| 1135 | sAD | F | 76 | AD | V (B3) | C3 |  |  | Nanostring |  |
| 461 | sAD | F | 96 | AD | VI (B3) | C3 | LB, amygdala predominant |  | Nanostring |  |
| 5805 | sAD | F | 64 | AD | VI (B3) | C3 |  |  | Nanostring |  |
| 220 | sAD | F | 83 | AD | V (B3) | C3 |  |  | Nanostring |  |
| 226 | sAD | F | 77 | AD | V (B3) | C3 | LB, brainstem predominant |  | Nanostring |  |
| 5986 | sAD | F | 64 | AD | VI (B3) | C2 |  |  | Nanostring |  |
| 1222 | sAD | M | 65 | AD | VI (B3) | C2 | LB, neocortical (diffuse) |  | Nanostring |  |
| 600 | sAD | F | 91 | AD | VI (B3) | C3 |  |  | Nanostring |  |
| 3087 | sAD | M | 68 | AD | VI (B3) | C3 |  |  | Nanostring |  |
| NHD 1 | PLOSL | M | 50 | PLOSL | NA | NA |  | TREM2 D134G | Nanostring | (Paloneva et al., 2002) |
| NHD 2 | PLOSL | F | 48 | PLOSL | NA | NA |  | TYROBP c.141delG | Nanostring | (Satoh et al., 2014) |
| NHD 5 | PLOSL | M | 38 | PLOSL | NA | NA |  | TYROBP c.141delG | Nanostring | (Satoh et al., 2014) |
| NHD 7 | PLOSL | M | 39 | PLOSL | NA | NA |  | TREM2 c.482+2T>C | Nanostring | (Numasawa et al., 2011) |
| 752 | CNT | M | 87 | CNT | 0 (B0) | C0 |  |  | RNA-seq |  |
| 2060 | CNT | F | 91 | CNT | II (B1) | C1 |  |  | RNA-seq |  |
| 750 | CNT | F | 93 | CNT | I (B1) | C0 |  |  | RNA-seq |  |
| 431 | CNT | M | 86 | CNT | II (B1) | C0 |  |  | Nanostring |  |
| 1845 | CNT | F | 73 | CNT | II (B1) | C0 |  |  | Nanostring |  |
| 420 | CNT | F | 87 | CNT | II (B1) | C0 |  |  | Nanostring |  |
| 1308 | CNT | F | 80 | CNT | III (B2) | C2 |  |  | Nanostring |  |
| 6049 | CNT | M | 56 | CNT |  |  |  |  | Nanostring |  |
| 1988 | CNT | F | 76 | CNT | II (B1) | C0 |  |  | Nanostring |  |
| 591 | CNT | F | 89 | CNT | III (B2) | C1 |  |  | Nanostring |  |
| 1362 | CNT | M | 72 | CNT | 0 (B0) | C0 |  |  | Nanostring |  |
| 310 | CNT | M | 87 | CNT | I (B1) | C1 |  |  | Nanostring |  |
| 1854 | CNT | F | 77 | CNT | III (B2) | C1 |  |  | Nanostring |  |
| 2040 | CNT | F | 77 | CNT | III (B2) | C0 |  |  | Nanostring |  |
| 1321 | CNT | F | 77 | CNT | I (B1) | C1 |  |  | Nanostring |  |
| 1389 | CNT | M | 71 | CNT | II (B1) | C0 |  |  | Nanostring |  |
| 2044 | CNT | F | 94 | CNT | II (B1) | C0 |  |  | Nanostring |  |
| 750 | CNT | F | 93 | CNT | I (B1) | C0 |  |  | Nanostring |  |
| 2599 | CNT | F | 71 | CNT | I (B1) | C0 |  |  | Nanostring |  |
| 224 | CNT | F | 70 | CNT | 0 (B0) | C0 |  |  | Nanostring |  |

Abbreviations:
M - male
F - female
LB - Lewy Bodies
PD - Parkinson's disease
FTLD - Frontotemporal lobar degeneration
NA - not applicable

**Table S2.** Neuropathologic, demographic and RNA QC characteristics of the groups for Nanostring gene expression analysis.

|  | CNT (N=17) | sAD (N=15) | TREM2 R47H AD (N=8) | PLOSL (N=4) |
| --- | --- | --- | --- | --- |
| Braak: AVE (min-max) | 1.8 (0-3) | 5.7 (5-6) | 5.5 (5-6) | NA |
| CERAD: AVE (min-max) | 0.4 (0-2) | 2.8 (2-3) | 2.3 (1-3) | NA |
| AGE, years: Ave (min-max) | 79 (56-94) | 74 (62-96) | 74 (64-89) | 42 (38-48) |
| Gender: males, % | 29,4% | 27% | 25% | 75% |
| PMI, hours: AVE (min-max) | 10 (2.5-22.2) | 12 (2.5-36) | 13 (7.6-18) | ND |
| RIN: AVE (min-max) | 2.5 (1.6 - 4.5) | 2.4 (2.1-3.1) | 2.4 (2-2.8) | 2.4 (2.2-2.6) |

NA – not applicable
ND – not determined

**Figure S1.**
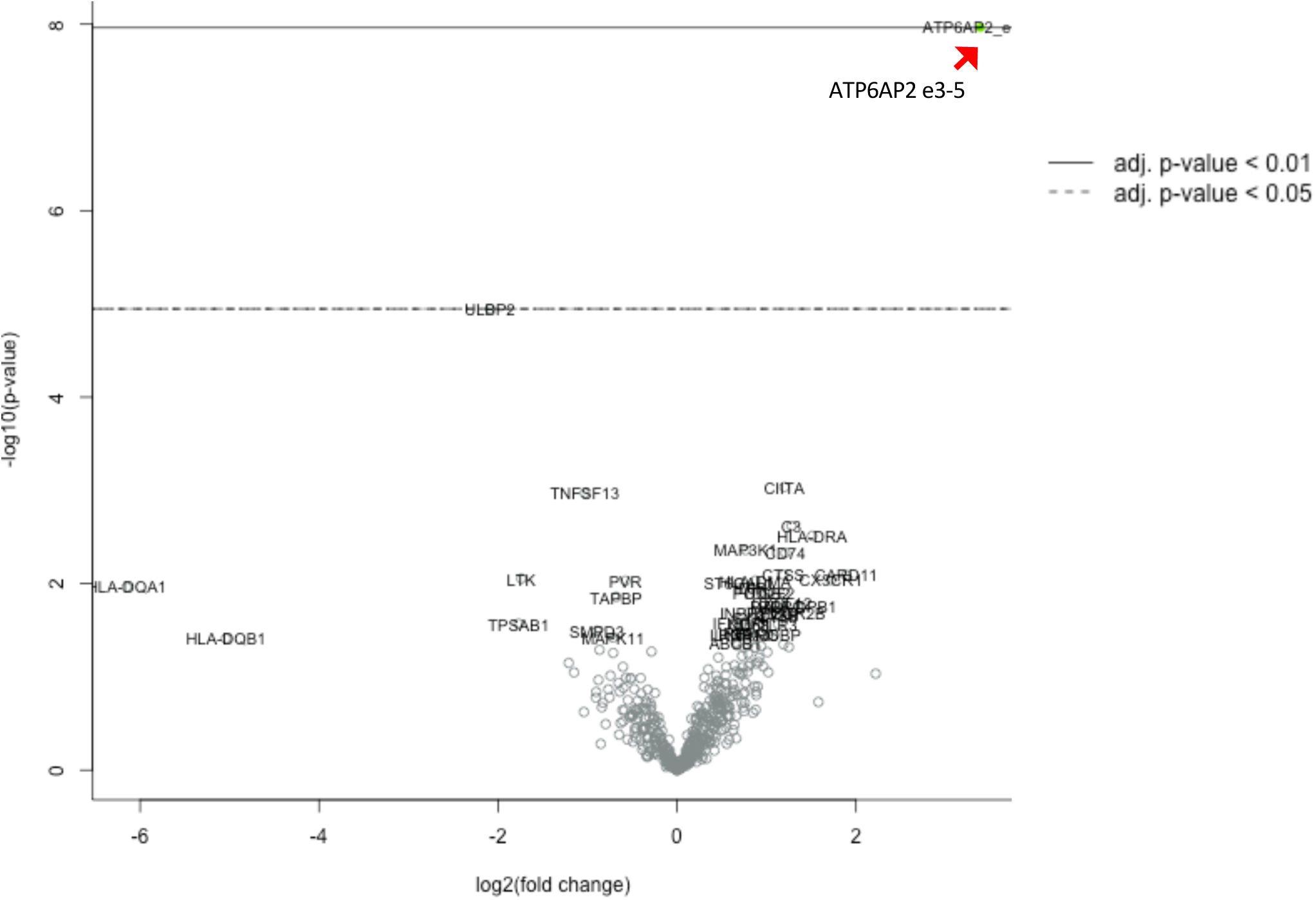
Nanostring nCounter detects upregulated abnormal ATP6AP2 e3-e5 splice isoform in RNA samples isolated from hippocampal FFPE sections of the ATP6AP2 mutation carriers. Volcano plot displaying DE genes in carriers (N=2) vs a baseline of gene expression in controls (N=17). Red arrow depicts position of ATP6AP2_e3–e5 splice-isoform. The y-axis corresponds to the log_10_(p-value), and the x-axis displays the log_2_(fold change) value. Carriers: female, 90 years; PMI: 25h, RIN :4.6; male, 86 years; PMI: ND; RIN: 1.8 Control subjects are listed in Tables S1, S2. A probe to e3-e5 junction of the *ATP6AP2* gene was designed as a part of PanelPlus Code Set and analyzed with a pre-designed Pan-Cancer Immune Panel.

**Figure S2.**
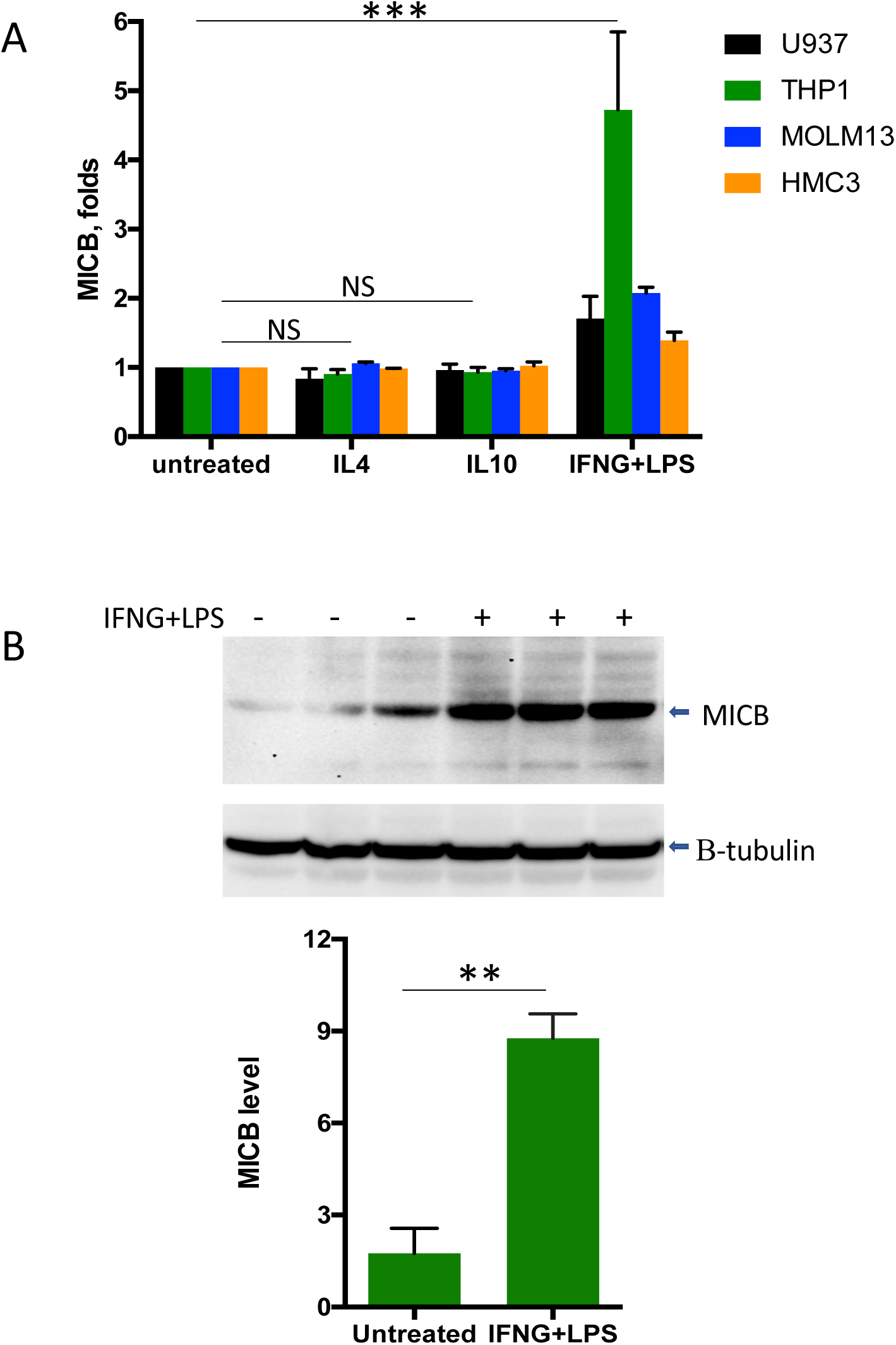
Pro-inflammatory stimulation leads to up-regulation of stress-ligand MICB in microglial (HMC3) and blood (THP1, MOLM13, U937) myeloid cell lines. (A). MICB mRNA expression was measured by qRT-PCR in cells treated with of IL4, IL10 or a combination of LPS with IFN−γ for 24h. MICB expression was normalized to its level in untreated cells. Data presented as mean ± SEM (*** - p-value < 0.001, two-way ANOVA, Dunnett’s multiple comparison test) (B). MICB protein expression in THP1 untreated or treated with LPS and IFN−γ for 24h was measured by Western blotting and normalized to Beta-tubulin. Data presented as mean ± SEM (** - p-value < 0.01, paired two-tailed T-test)

**Figure S3.**
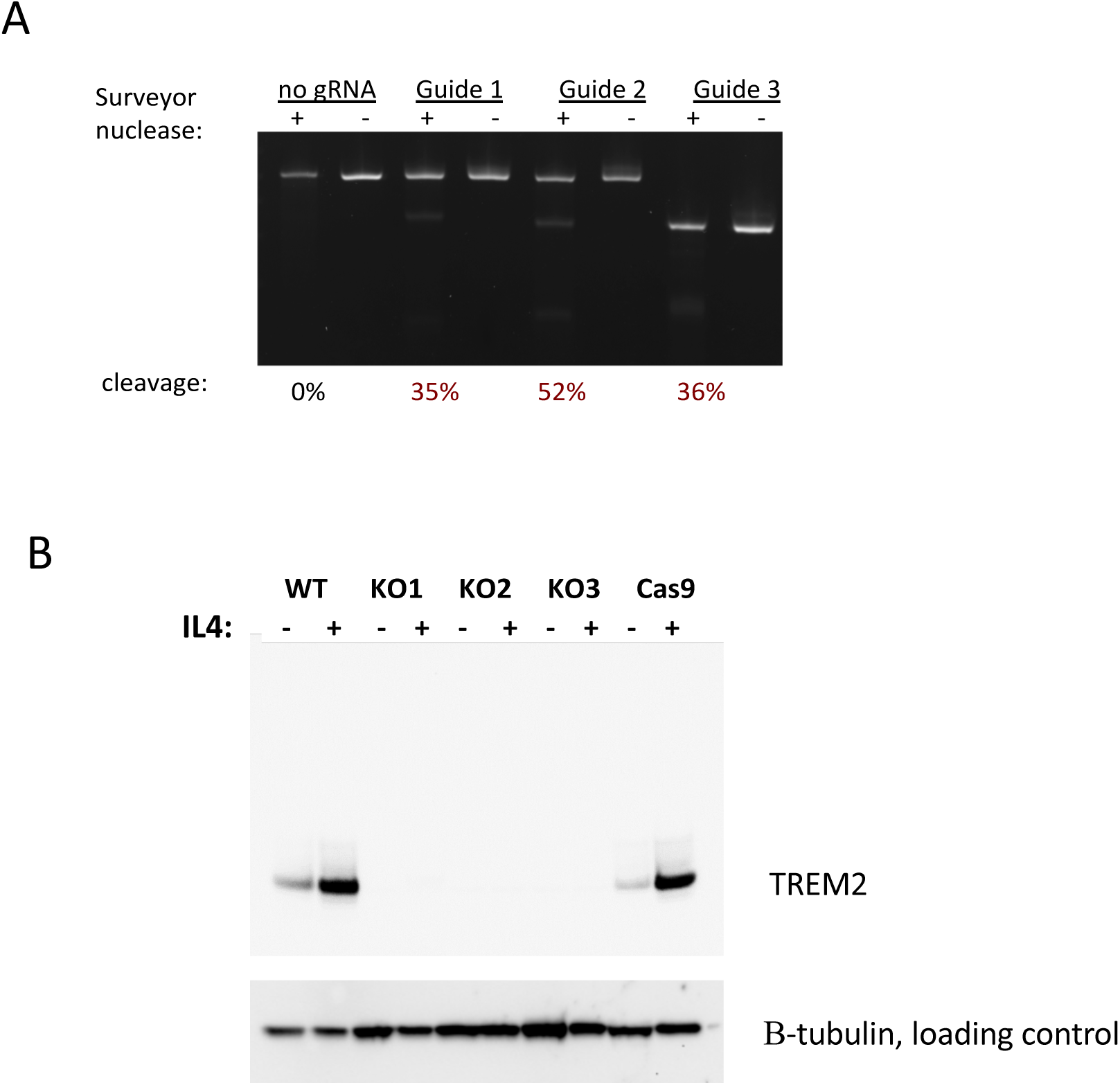
CRISPR/Cas9-mediated genome editing in THP-1 cells. (A) Efficiency of cleavage by different guide RNAs evaluated by Surveyor nuclease assay. (B) Expression of TREM2 protein in THP-1 (WT), cells stably expressing Cas9 nuclease and three independent knockout clones of TREM2 (KO1-3). TREM2 level in cell extracts was measured by Western blot. TREM2 expression was stimulated with IL4.

